# Nuclear hormone receptor regulation of PAL-1/Caudal mediates ventral nerve cord assembly in *C. elegans*

**DOI:** 10.1101/2025.03.02.641102

**Authors:** Nathaniel Noblett, Tony Roenspies, Stephane Flibotte, Antonio Colavita

**Affiliations:** Neuroscience Program, Ottawa Hospital Research Institute, Ottawa, Canada; Department of Cellular and Molecular Medicine, University of Ottawa, Ottawa, Canada; UBC/LSI Bioinformatics Facility, University of British Columbia, Vancouver, Canada

**Keywords:** *C. elegans*, motor neuron positioning, ventral nerve cord, *Caudal*, PAL-1, SEX-1

## Abstract

The regulatory network governed by CDX/*Caudal* family transcription factors plays critical roles in shaping embryonic neural development. In *C. elegans*, we found that proper expression of *pal-1*, the *C. elegans Caudal* homologue, is required for correct positioning of motor neuron cell bodies in the first larval stage ventral nerve cord (VNC). We identified an upstream regulatory region within the *pal-1* promoter that drives *pal-1* expression in a subset of DD and DA neuronal progenitors. We also show that SEX-1, a nuclear hormone receptor, is required for motor neuron positioning in the VNC. Loss of *sex-1* results in neuronal positioning defects similar to those observed in *pal-1* mutants. This is in part due to a requirement for SEX-1 in promoting *pal-1* expression in DD and DA neuronal progenitors during VNC assembly. Double mutant analysis further suggests that *sex-1* also has *pal-1*-independent functions. Together, these findings define a transcriptional hierarchy in which the SEX-1 nuclear hormone receptor regulates the tissue-specific activity of PAL-1 to promote proper motor neuron positioning in the VNC and highlight a conserved role for NHR and CDX/Caudal family proteins in central nerve cord formation.

**Highlights:**

- PAL-1 is required for proper neuron cell body positioning in the ventral nerve cord (VNC) in newly hatched worms.
- An upstream promoter element controls the expression of *pal-1* in DD and DA neurons.
- The nuclear hormone receptor SEX-1 is required for proper neuron positioning in the VNC.
- SEX-1 regulates PAL-1 expression in DD and DA neurons.

## Introduction

The coordination of signaling cascades and positional information through transcriptional networks is essential for axial extension during embryonic morphogenesis (Collinet and Lecuit, 2021). The *Caudal* family of homeodomain transcription factors, including CDX1/2/4 in mammals and their worm homologue PAL-1, play critical roles in posterior patterning and axial extension by coordinating pathways involved in cell-fate determination, morphogenesis and regionalization (Edgar et al., 2001; van den Akker et al., 2002; Gilbert et al., 2020; Zhu and Lohnes, 2022). For CDX proteins this includes the regulation of neurulation during mammalian development, a process where the neuroectoderm folds and elongates to form the neural tube (Savory et al., 2011; Zhao et al., 2014). Neurulation is partly driven by convergent extension, a morphogenetic process characterized by coordinated cell-cell intercalations along the mediolateral axis and extension along the anterior-posterior axis. Mouse embryos with mutations in both CDX1 and CDX2 exhibit typical convergent extension defects, including a reduced length-to-width ratio of the neural tube, failure of extension, and a broadened midline of the neural plate (Savory et al., 2011). These defects lead to embryonic lethality and craniorachischisis, a severe neural tube defect (Savory et al., 2011).

CDX proteins function as a central hub for multiple signaling pathways that regulate neurulation (Wilson et al., 2003; Palmer et al., 2021; Zhao et al., 2022). This includes the direct induction of CDX1 by retinoic acid receptors (RARs) α1 and γ, two types of ligand dependent nuclear hormone receptors, as well as β-catenin, a key component of canonical WNT signaling (Houle et al., 2003; Zhao et al., 2014). During neurulation, CDX members have been linked to regulation of the Planar Cell Polarity (PCP) pathway, a key pathway involved in convergent extension (Savory et al., 2011). PCP components associated with CDX include Scribble and the transmembrane receptor PTK7 (Wehner et al., 2011; Hayes et al., 2013; Martinez et al., 2015). Both CHiP analysis and transfection assays suggest that CDX proteins directly regulate PTK7 expression (Savory et al., 2011).

In *C. elegans*, PAL-1 is involved in cell fate determination, regional identity, and morphogenesis. PAL-1 plays an early role in blastomere cell fate, including a key role in establishing the fate of descendants of the C and D blastomeres (producing primarily body-wall muscle, hypodermal cells, along with two neurons) and, along with the WNT effector POP-1 and the bZIP transcription factor SKN-1, the descendants of the E blastomere (producing intestinal cells) (Hunter and Kenyon, 1996; Maduro et al., 2005). Additionally, PAL-1 is involved in non-cell fate related functions, such as orienting epidermal seam-cell migration and alignment during development (Gilbert et al., 2020). Other defects, including delays in ventral enclosure, failed long range migration in MS cell descendants, and abnormal cleavage axis in muscle cells suggest that additional functions of a *pal-1* network remain to be explored (Edgar et al., 2001).

At hatching, the ventral nerve cord (VNC) in *C. elegans*, is comprised of 22 embryonically born motor neurons, belonging to three classes (6 GABAergic inhibitory DD, 9 cholinergic excitatory DA, and 7 cholinergic excitatory DB), that innervate body wall muscle to control sinusoidal locomotion (Lu et al., 2022). Progenitors of these neurons are born on the left and right sides of the embryo and undergo collective cell movement toward the midline where they intercalate to form the VNC. This process shares several key features with neurulation in mammals, including rosette mediated-convergent extension, asymmetric distribution of core PCP components like VANG-1/VangL, and inositol phosphate signaling (Shah et al., 2017; Noblett et al., 2025).

Here, we describe roles for the nuclear hormone receptor (NHR) SEX-1 and the *Caudal* homologue PAL-1 in regulating proper neuronal organization of the VNC. Using alleles with tissue-specific effects on gene expression, we identified a role for *pal-1* in regulating the proper positioning of embryonically derived motor neurons in the VNC. These mutations disrupt a *pal-1* promoter region required for expression in a subset of neuronal progenitors during embryonic convergent extension movements that form the VNC. We found that SEX-1 is required for *pal-1* expression in progenitor neurons, and that restoring *pal-1* expression in these cells is sufficient to partially rescue the neuronal positioning defects in *sex-1* mutants. These findings show that nuclear hormone regulation of Caudal is important for VNC assembly. Given that NHRs and CDX/Caudal proteins contribute to neural tube formation in vertebrates (Barreto et al., 2003; Savory et al., 2011; Zhao et al., 2014; Koch et al., 2025), our findings highlight the deep conservation of mechanisms underlying central nerve cord development.

## Results

### Identification of a *pal-1* promoter region involved in neuronal specific expression

The VNC at the first larval (L1) stage contains three classes of motor neurons (6 DD, 9 DA, and 7 DB). We have previously shown that loss of PCP genes, such as *vang-1*, and the Robo encoding gene *sax-3* result in VNC assembly defects during embryogenesis, which manifest as mispositioned motor neurons in L1 larvae (Shah et al., 2017). To identify new genes involved in neuron positioning, we performed a genetic screen using the DD-specific reporter *ynIs37[flp-13p::GFP]* (unpublished results). We identified two mutants, *zy43* and *zy44*, that displayed similar DD neuron positioning defects, with DD2 and DD3 positioned closer together than in wild type (WT) (Fig. 1A). Whole genome sequencing revealed these mutations to be single nucleotide changes, one nucleotide apart, located 143 and 145 bp upstream of the *arl-6* start codon (Fig. 1C). To determine if these changes were responsible for the neuron position defects, we used Cas9-targetted non-homologous end joining-mediated DNA repair to generate a 44 bp deletion (*zy117*) centered on the mutated region (Fig. 1C and D). This mutant displayed similar neuron positioning defects (Fig. 1B).

**Figure 1.**
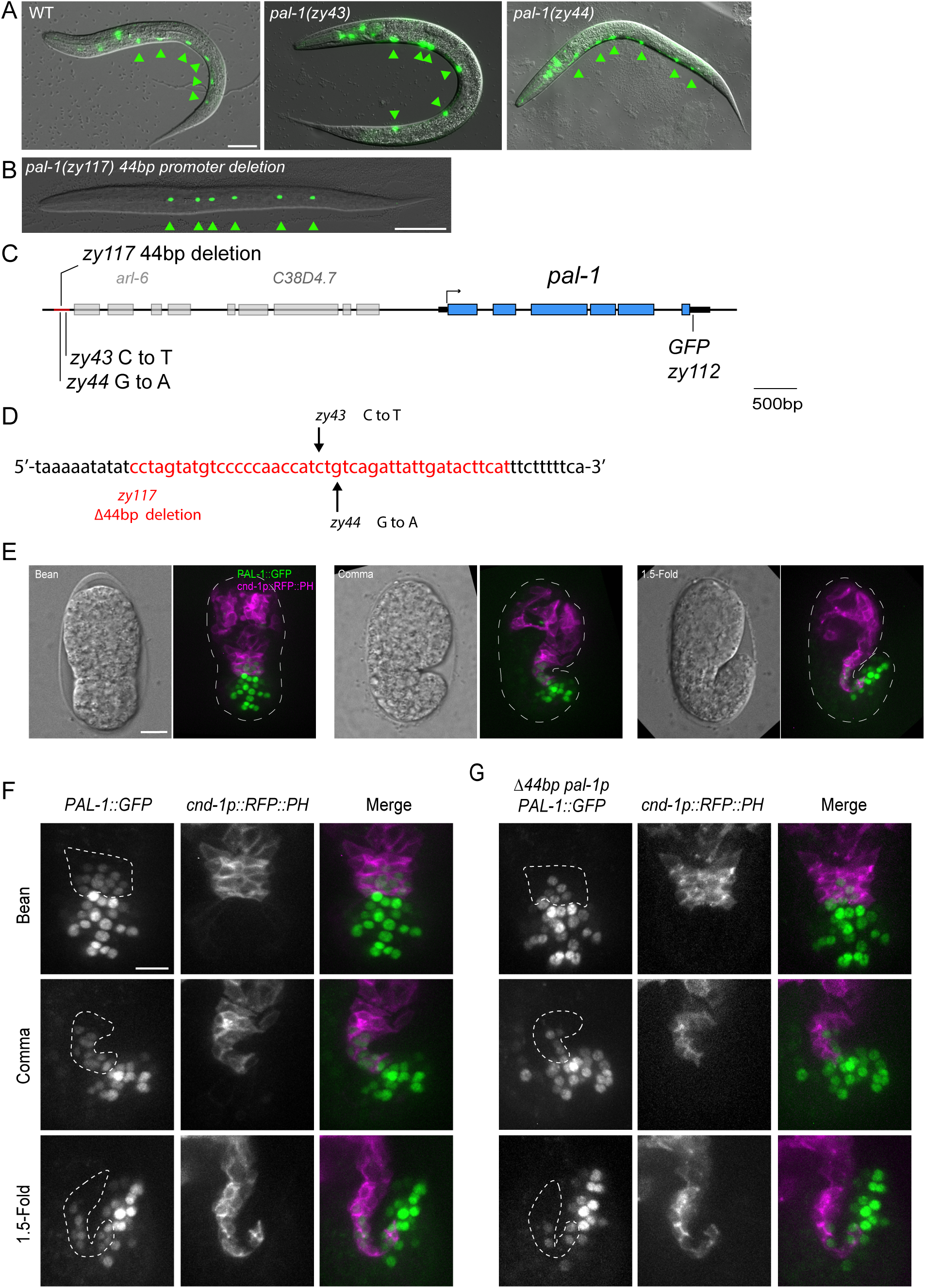
A distal promoter element regulates *pal-1* expression in DD and DA progenitors. (A-B) Representative images of DD positions in (A) WT, *pal-1(zy43)*, *pal-1(zy44)* and (B) *pal-1(zy117)* mutants. Arrowheads mark DD neurons, which are visualised using *flp-13p::GFP*. Scale bar = 20μm. (C) Diagram of the *pal-1* genomic region showing the positions of mutations used in this study. (D) The location of nucleotide changes in *zy43*, *zy44* and *zy117*. Red nucleotides highlight the deleted sequence in *pal-1(zy117)*. (E) Representative maximum intensity projections of an endogenous *pal-1::GFP* expression at three stages of embryonic development. Nomarski image indicates the morphology of the embryo. A *cnd-1p::mCherry ::PH* reporter labels the membranes of DD and a subset of DA progenitors. Scale bar = 10μm. (F) Representative maximum intensity projections show endogenous *pal-1::GFP* expression in WT and *pal-1(zy117)* mutant backgrounds. The dotted line indicates regions of interest containing expression changes. Number of embryos examined: N = 8 for WT and N = 3 for *pal-1(zy117)* at the bean stage. N = 4 for both genotypes at comma stage. N = 7 for WT and N = 6 for *pal-1(zy117)* at the 1.5-fold stage. Scale bar = 10μm.

The proximity of *zy43* and *zy44* to the *arl-6* gene led us to examine DD neuron positioning in an *arl-6* mutant, but we did not observe any defects (data not shown). These mutations were also located approximately 6.5 kb upstream of *pal-1* (Fig. 1C), an essential gene encoding an homologue of the homeobox protein CDX/Caudal, within a region known to be required for complete rescue of *pal-1* embryonic defects (Edgar et al., 2001). The CDX family of proteins are also associated with neural tube defects in mammals, suggesting *pal-1* was a good candidate for involvement in VNC formation (Savory et al., 2011).

Because strong *pal-1* loss-of-function mutants are non-viable and display severe embryonic and larval morphology defects (Baugh et al., 2005), we asked whether the 44 bp *zy117* deletion disrupted a promoter region important for *pal-1* expression. To address this, we endogenously tagged the *pal-1* gene with a C-terminal GFP in both the WT and *zy117* background (Fig. 1E). As previously reported, PAL-1::GFP is widely expressed in the nuclei of posterior cells from the bean to 1.5-fold stage embryo (Edgar et al., 2001). Using a *cnd-1p::mCherry::PH* reporter (*zyIs36*) to label the membranes of DD and DA neuronal progenitors, we observed that *PAL-1::GFP* was also expressed in the nuclei of a subset of DD, DA, and DB progenitors, specifically those located in the posterior VNC (Fig. 1E, F). Based on their location, these appear to be DD3-6, DA3-6, and DB5-7. Strikingly, in the *zy117* promoter mutant, PAL-1::GFP expression was completely absent from DD and DA progenitors (Fig. 1G). Posterior tail cells and DB progenitors, which can be identified by their location outside and ventral to *cnd-1p*-labeled progenitors at the 1.5-fold stage, were not affected in *zy117* mutants. These results indicate that *zy43*, *zy44*, and the 44 bp *zy117* deletion mutant disrupt a *pal-1* promoter regulatory element required for expression in DD and DA neurons.

### *pal-1* promoter mutants exhibit motor neuron cell body positioning defects

Our *pal-1* promoter mutants were identified based on defects in the positioning of DD motor neurons, one of three classes of motor neurons present in the VNC of newly hatched L1 worms. To quantify these position defects and determine if they extend to the other two classes of motor neurons, DA and DB, we utilised a two-colour reporter background in which each neuron class was uniquely labeled with a class-specific fluorescent cell fate marker at L1 (Saharkhiz et al., 2024). The nuclei of DD1-6 were labeled in red with *zySi2[unc-30p::mCherry::H2B]*, DA1-9 in green with *unc-4(zy123[unc-4::mNG])*, and DB1-7 in both red and green, detected as yellow, with *vab-7(zy142[vab-7::mNG::T2A::mScarlet-I::H2B])*. The *unc-4* marker also labels the interneurons SABVL, SABVR, and SABD, which are located anterior to DA1. Neuron positions were normalized by setting SABVL as the anterior reference point (0%) and the rectum as the posterior reference point (100%), allowing neuron locations to be compared relative to VNC length across different animals. When plotted, neurons of each class are numbered sequentially from anterior to posterior, regardless of their terminal cell lineage identity, which may be unknown in mutant backgrounds.

DD, DA, and DB motor neurons are stereotypically positioned and arranged in a repeating pattern along the VNC. We found that the mean position of most DD neurons in *pal-1(zy117)* showed significant anterior shifts along the VNC compared to WT (Fig. 2A and B, Fig. S1). Some DA and DB neurons were also mispositioned compared to WT, but these shifts were milder than the DD neuron defects and may represent secondary effects caused by the mispositioned DD neurons. Indeed, the strongest position shifts affected DD3–6, which correlates with *pal-1* expression in these neurons, while milder shifts were observed in DD1 and DD2, where expression is absent. In addition, except for DA8 and DA9, which are located opposite each other on the left and right in the preanal ganglia, the neuron classes are arranged in an alternating sequence, with no two classes adjacent to each other (Fig. 2C-E). This pattern is significantly altered in *zy117* mutants, with some same class cell bodies more frequently adjacent to each other (Fig. 2D and E). For example, in 21% of *zy117* larvae (N=53), the DD2 and DD3 cell bodies are adjacent with no intervening cells, compared to 0% in WT (N=51). Notably, although *pal-1* regulates cell fates (Edgar et al., 2001; Maduro et al., 2005), we did not observe any changes in DD, DA, or DB cell numbers that would indicate cell fate defects during our analysis. Overall, these results indicate that *pal-1* expression, driven by the DD-DA-specific promoter element, is primarily required for proper DD motor neuron positioning.

**Figure 2.**
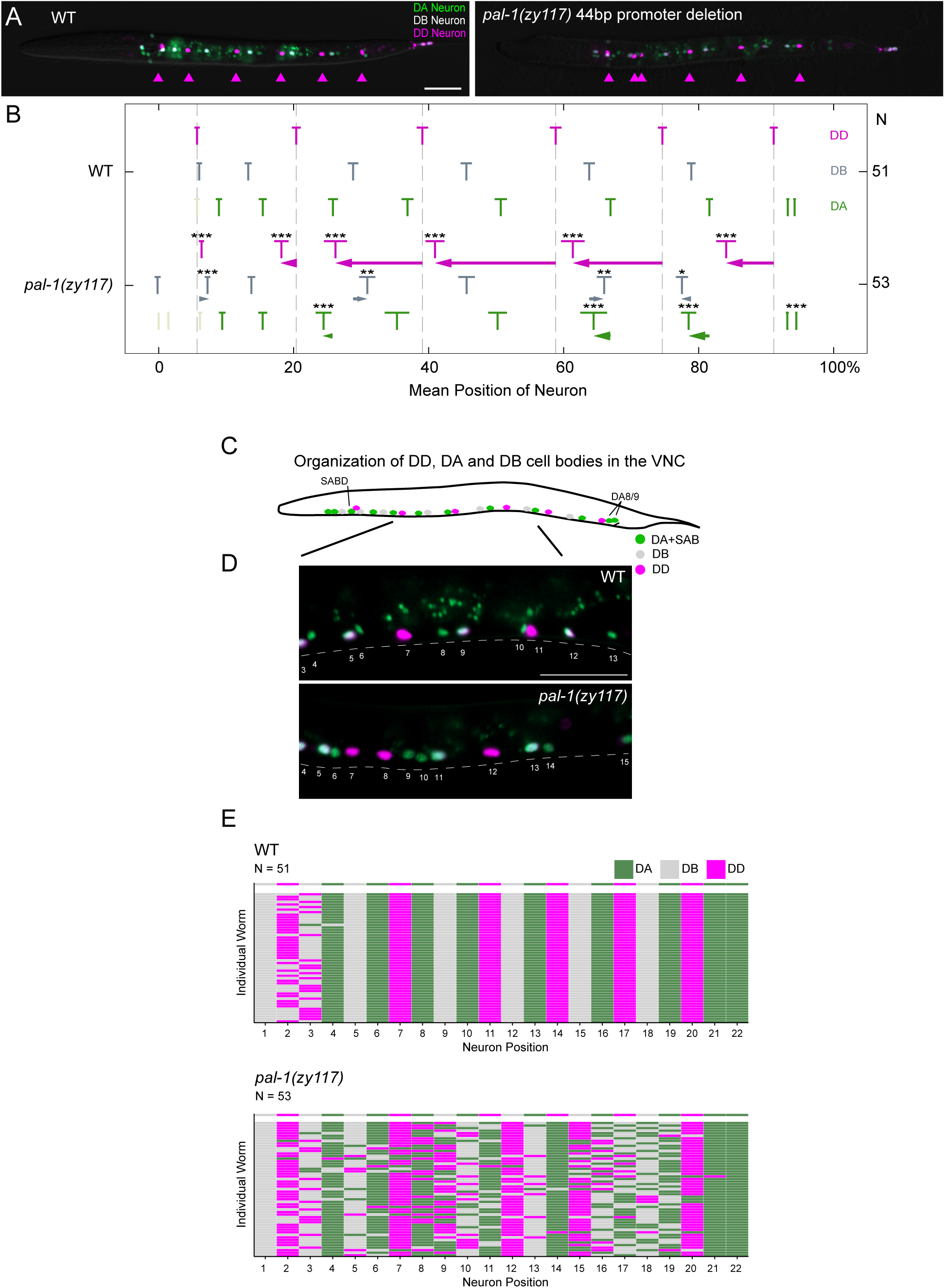
*pal-1* is required for proper motor neuron positioning in the VNC. (A) Representative images of DD, DA, and DB positions in WT and *pal-1(zy117)* mutants. DA neurons (*zy123[unc-4::mNG*) are indicated in green, DB neurons (*zy142[vab-7::mNG::T2A::mScarlet-I::H2B]*) in grey and DD neurons (*unc-30p::mCherry::H2B*) in magenta. Arrowheads mark the position of DD neurons. Scale bar = 20μm. (B) Quantification of DD, DA, and DB mean neuron position relative to the SABVL neuron and the rectum in L1 stage worms. Means and 95% confidence intervals are indicated for each neuron. Animals scored (N) indicated on the right. Hashed lines indicate the mean position of WT DD neurons. Arrows indicate the size and direction of the mean neuron position in *pal-1(zy117)* from the mean position of their WT equivalent. (C) Schematic of neuron organization in the L1 stage VNC. DA neurons in green and DD neurons in magenta. (D) Representative images showing neuron organization in WT and *pal-1(zy117)* mutants. Scale bar = 20μm. (E) Organization of DA, DB and DD neurons in WT and *pal-1(zy117)* worms. Individual boxes show the identity of the neuron in each position along the VNC with each row indicating an individual. The top row shows the stereotypical order of neurons in WT. Statistics: (B) Weltch’s T-Test. *p<0.05, **p<0.01, ***p<0.001.

### PAL-1 acts independently of VANG-1 and SAX-3 to position motor neurons in the VNC

We previously showed that the PCP pathway component VANG-1 and the Robo receptor SAX-3 act in parallel to regulate motor neuron cell body positioning in the VNC of newly hatched larvae. Loss of both *vang-1* and *sax-3* causes a highly penetrant anterior displacement of motor neuron cell bodies, a defect that is synergistically more severe than in either single mutant (Shah et al., 2017). This pronounced anterior shift reflects a major disruption of convergent extension movements among tightly juxtaposed neuronal progenitors during embryogenesis. From their birth locations on the left and right sides of the embryo, these progenitors converge mediolaterally, narrowing the tissue along the medial-lateral axis while elongating the nascent VNC along the anterior-posterior axis (Shah et al., 2017). In *vang-1 sax-3* double mutants, elongation of VNC progenitors along the anterior-posterior axis is impaired or delayed, leading to neuronal cell bodies that are positioned significantly more anteriorly than in WT.

To investigate genetic interactions between *pal-1* and these pathways, we examined the VNC positions of DD and DA neurons relative to SABVL and the rectum in *pal-1(zy117)* double mutants with *vang-1(tm1422)* and *sax-3(zy5)* (Fig. 3A). *tm1422* is a deletion allele predicted to eliminate VANG-1 activity, whereas *zy5* is a nonsense allele expected to produce a SAX-3 protein with a truncated cytoplasmic domain (Shah et al., 2017). The *zy5* allele is used instead of the *sax-3* null allele because it exhibits significantly higher viability and fewer body morphology defects, which would otherwise interfere with measurements of neuron position. We found that the anterior shifts in the mean position of DD3-6 cell bodies along the VNC in *pal-1(zy117)* mutants were substantially more severe than those in *vang-1* and *sax-3* single mutants (Fig. 3B and Fig. S2). In contrast, the anterior shifts in DA cell bodies were similar across all mutants, being relatively mild in each case. *vang-1; pal-1* double mutants exhibited significantly more severe anterior shifts in mean DD and DA neuron positions compared to either single mutant. Interestingly, in *sax-3; pal-1* double mutants, the mean position of DD cell bodies showed significantly more severe anterior shifts compared to single mutants, while only DA3 and DA4 among the DA cell bodies were significantly different from *pal-1* mutants (Fig. 3B and Fig. S3). These results suggest that, at least for some aspects, *pal-1* and *vang-1* act in parallel or independent pathways to position DD and DA cell bodies. This redundancy is particularly evident in DA neurons which show a synergistic increase in the anterior shifts of mean cell body position compared to the single mutants. The findings also suggest that *pal-1* and *sax-3* act in parallel or independent pathways to position DD neurons and a subset of DA neurons. Furthermore, the similarity of the *vang-1; pal-1* cell body position phenotype to the *vang-1; sax-3* phenotype, in which convergent extension is disrupted, suggests that the downstream transcriptional targets of *pal-1* may also act to mediate convergent extension.

**Figure 3.**
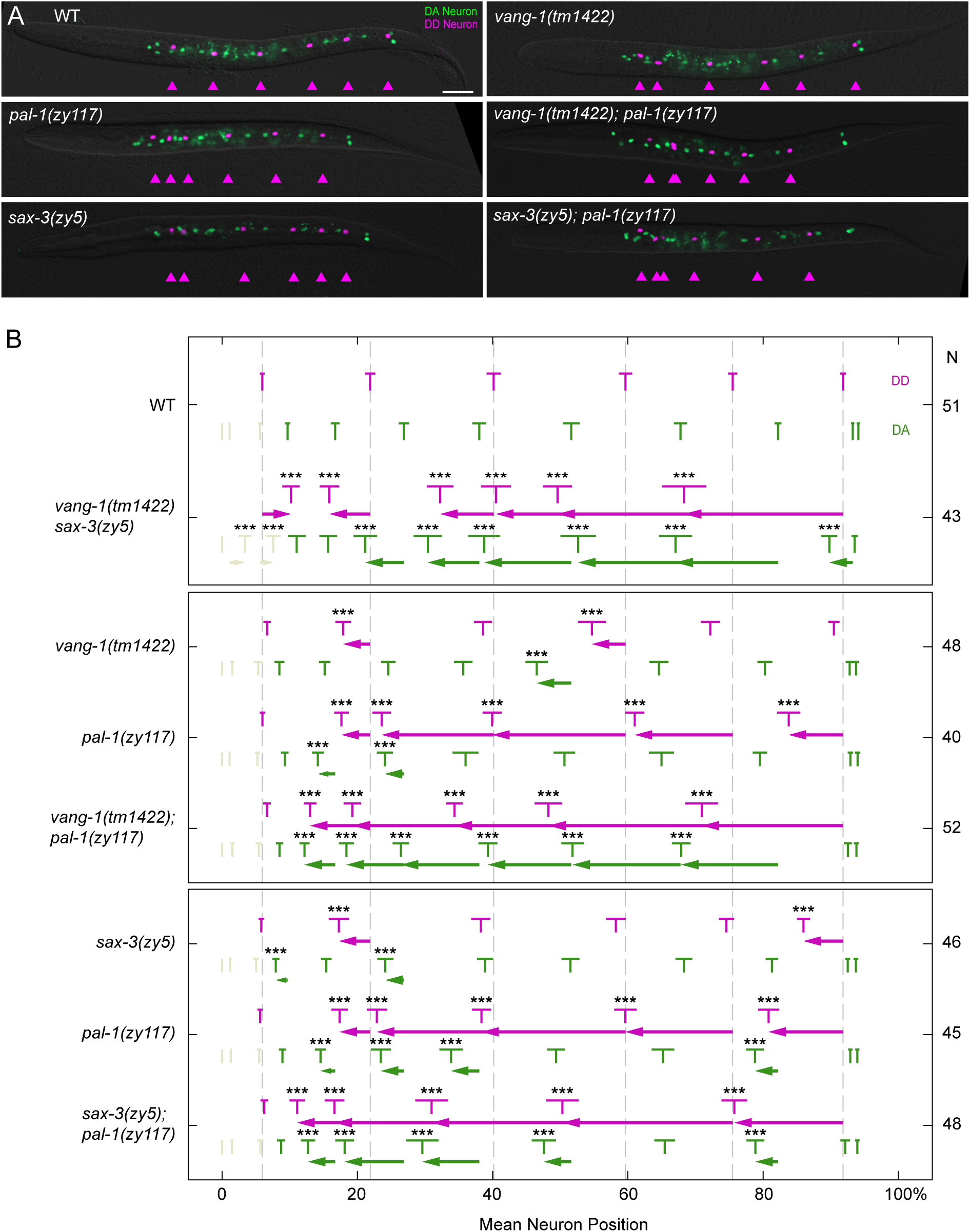
Genetic interactions between *pal-1*, *vang-1*, and *sax-3* during neuronal positioning. (A) Representative images of WT, *pal-1(zy117), vang-1(tm1422)*, *sax-3(zy5)* single and double mutants. DA neurons in green and DD neurons in magenta. Arrowheads mark the position of DD neurons. Scale bar = 20μm. (B) Quantification of DD and DA neuron position relative to the SABVL neuron and rectum in L1 stage worms. Means and 95% confidence intervals are indicated for each neuron and animals scored (N) indicated on the right. Hashed lines indicate the position of WT DD neurons. Arrows indicate the size and direction of the mean neuron position shift in each neuron compared to the mean position of their WT equivalent. Statistics: (B) one-way ANOVA with Tukey’s Post-hoc Comparison. *p<0.05, **p<0.01, ***p<0.001.

### *sex-1* mutants exhibit motor neuron cell body positioning defects

*sex-1* encodes a 534-amino acid nuclear hormone receptor that was originally identified for its key role in sex determination (Carmi et al., 1998). In the same screen that identified *pal-1 zy43* and *zy44*, we also isolated a *sex-1* mutant. This mutant exhibited abnormally close spacing between the DD2 and DD3 cell bodies, similar to the defects observed in *pal-1* mutants (Fig. 4A). This mutation, *sex-1(zy45)*, introduces a premature stop codon at amino acid W51, predicted to result in a severely truncated protein. We confirmed that *sex-1* is the causative gene by observing similar motor neuron positioning defects in two additional CRISPR/Cas9-generated *sex-1* alleles: *zy130*, a 7-nucleotide deletion that causes a frameshift after L54 and is likely a null allele, and *zy159*, a 20-nucleotide deletion that causes a frameshift after V253. These alleles were originally created for an unrelated study (unpublished). Given that *pal-1* and *sex-1* show a similar DD2–DD3 cell body proximity phenotype, we investigated the role of *sex-1* in neuronal positioning.

**Figure 4.**
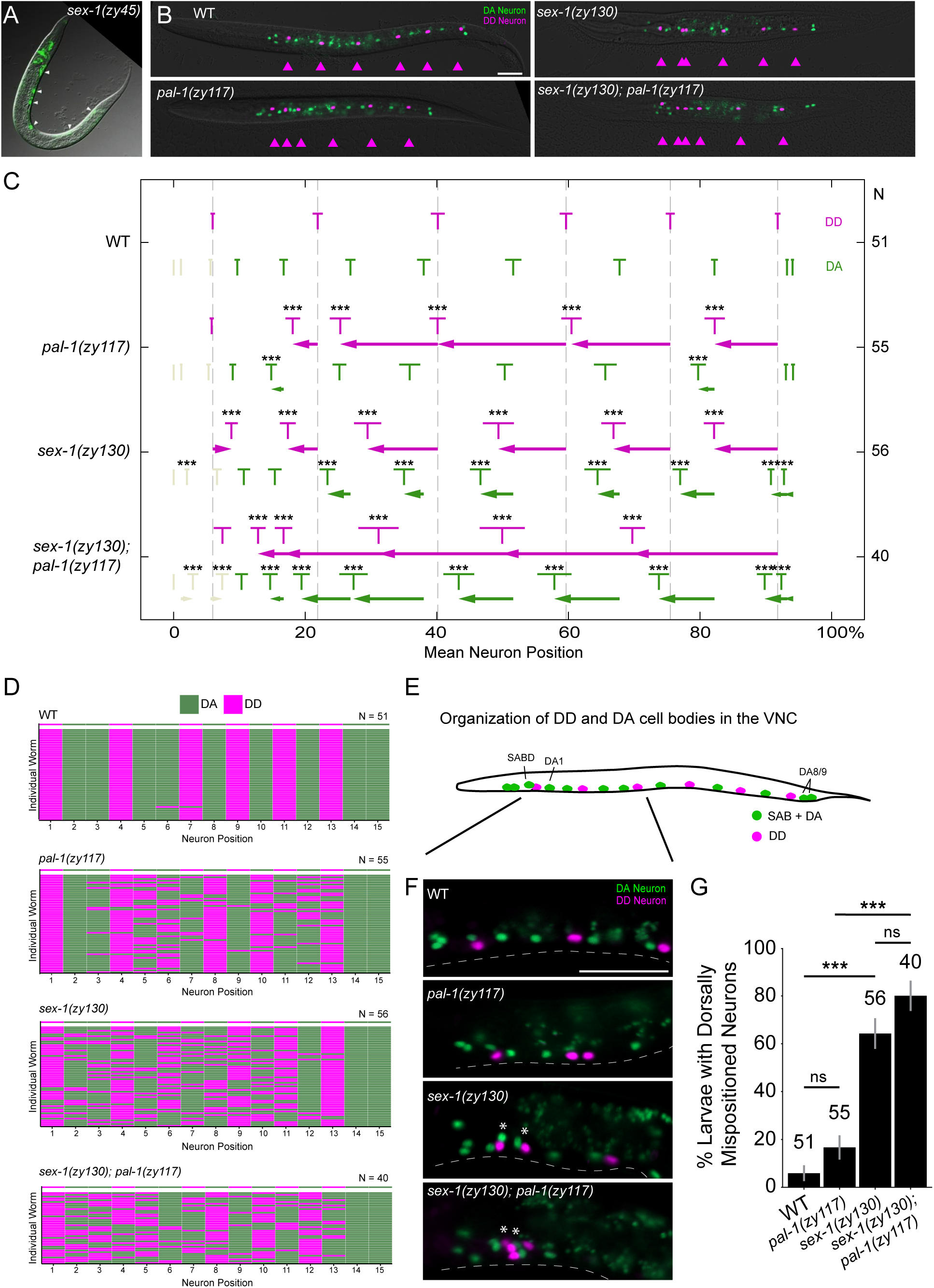
*sex-1* is required for proper motor neuron positioning in the VNC and genetic interactions with *pal-1*. (A) Representative image of DD neuron positions in *sex-1(zy45)*, visualized using *flp-13p::GFP*. (B) Representative images of WT, *pal-1(zy117)*, *sex-1(zy130)*, and double mutants showing DA neurons in green and DD neurons in magenta. Arrowheads mark the position of DD neurons. Scale bar = 20μm. (C) Quantification of DD and DA neuron position relative to the SABVL neuron and rectum in L1 stage worms. Means and 95% confidence intervals are indicated for each neuron. Animals scored (N) indicated on the right. Hashed lines indicate the position of WT DD neurons. Arrows indicate the size and direction of the mean neuron position shift in each neuron from the mean position of their WT equivalent. (D) Organization DD and DA neurons in WT and mutant worms. (E) Schematic showing DD and DA neuron organization along the VNC. (F) Representative images of anterior DA and DD neurons in *pal-1(zy117), sex-1(zy130)* and double mutants. Asterisks mark dorsally mispositioned neurons. Scale bar = 20μm. (G) Quantification of worms with dorsally mispositioned neurons at L1. Statistics: (C) one-way ANOVA with Tukey’s Post-hoc Comparisons, and (G) Chi-squared test with Monte Carlo simulation for 10,000 replicates, followed by a pairwise analysis using Fisher’s exact test with Monte Carlo simulation for 10,000 replicates and adjusted with Holm corrections. *p<0.05, **p<0.01, ***p<0.001.

To better understand motor neuron positioning defects in *sex-1(zy130)* null mutants, we used our neuron class-specific reporters to measure the positions of DD and DA cell bodies in the newly hatched VNC relative to SABVL and the rectum. *sex-1* mutants show significant anterior shifts in the positions of both DD and DA cell bodies compared to WT, in contrast to *pal-1(zy117)* mutants, which primarily affect DD neurons (Fig. 4B and C). These DD and DA cell body shifts become significantly more severe in *sex-1(zy130); pal-1(zy117)* double mutants compared to either single mutant (Fig. 4C and Fig. S4). As in *pal-1* mutants, cell body position shifts in *sex-1* mutants are associated with changes to the stereotypical arrangement of motor neurons along the VNC (Fig. 4D). However, unlike *pal-1* mutants, *sex-1* mutants also display additional organizational defects in the anterior VNC, where cell bodies are frequently mispositioned dorsally rather than aligned in a single file (Fig. 4E-G). The proportion of dorsally mispositioned neurons increased from 5.9% in WT to 64.3% in *sex-1(zy130)* mutants (p< 0.001) (Fig. 4G). Dorsally mispositioned neurons in *pal-1(zy117)* are not significantly different from WT (Fig. 4G). Mispositioning defects in *sex-1(zy130); pal-1(zy117)* double mutants appear additive but are not significantly more severe than those in *sex-1(zy130)* alone (Fig. 4G), consistent with *pal-1*, or at least its activity in DD and DA neurons, acting in a parallel pathway or not being involved in single file alignment. Together, these findings indicate that, at least for some aspects, *pal-1* and *sex-1* act redundantly or independently to position motor neurons in the VNC. Furthermore, *sex-1* plays an additional role in ensuring the single-file alignment of neurons in the anterior region of the VNC.

### SEX-1 regulates PAL-1 expression in DD and DA neuronal progenitors

Motor neuron positioning defects in the VNC of newly hatched worms indicate a disruption in the assembly of the VNC that begins at the embryonic bean stage. VNC assembly involves several distinct processes, including convergent extension of DD, DA, and DB neuronal progenitors, as well as still unknown sorting and spacing mechanisms that establish their stereotypical organization (Shah et al., 2017; Saharkhiz et al., 2024). The increased severity of neuron positioning defects in *sex-1; pal-1* double mutants compared to single mutants suggests that *sex-1* and *pal-1* may act in parallel or independent pathways during VNC assembly, though a role in the same pathway for some aspect of this process cannot be excluded. Given that both *sex-1* and *pal-1* encode transcription factors, we tested the possibility that one might regulate the expression of the other.

We first tested whether *pal-1* regulates *sex-1* expression. To do this, we knocked mNeonGreen (mNG) into the C-terminus of *sex-1* and examined its expression in WT and *pal-1(zy117)* mutant backgrounds, using our mCherry membrane reporter (*zyIs36*) to label DD and DA neurons. SEX-1::mNG was broadly expressed through the comma stage, including in the nuclei of all DD, DA, and DB progenitors (Fig. S5A). Its expression declined by the 1.5-fold stage, with few nuclei still showing detectable signal. No differences in SEX-1::mNG expression were observed in the *pal-1(zy117)* mutant background (Fig. S5B).

We next tested whether *sex-1* regulates *pal-1* expression. PAL-1::GFP is normally expressed and localized to the nuclei of a subset of DD, DA, and DB progenitors. In the *sex-1(zy130)* null background, PAL-1::GFP was absent from DD and DA progenitors but remained unaffected in DB progenitors at both the bean and 1.5-fold stages (Fig. 5A). To confirm these observations, we quantified PAL-1::GFP expression based on whether it was co-expressed with *cnd-1* (DD and DA) or not co-expressed (DB and tail cells). To do this, we used Imaris tools to segment PAL-1::GFP nuclear expression. These nuclei were then grouped based on whether they expressed *cnd-1* or not, and the mean sum intensity per cell was compared across genotypes. We found that the average intensity of PAL-1::GFP in *cnd-1* nuclei was dramatically reduced in embryos carrying the 44 bp *pal-1* promoter deletion and in both *sex-1* mutants (Fig. 5B). In contrast, *cnd-1* nuclei showed no significant change. This difference in intensity reflects the number of PAL-1::GFP positive nuclei detected in each region (Fig. 5C). We further showed that this regulation occurs at the level of *pal-1* transcription by showing reduced GFP expression in embryos carrying a transgene driven by the *pal-1* promoter (Fig. 5D). These findings indicate that PAL-1 expression in DD and DA neuronal progenitors requires transcriptional regulation by SEX-1.

**Figure 5.**
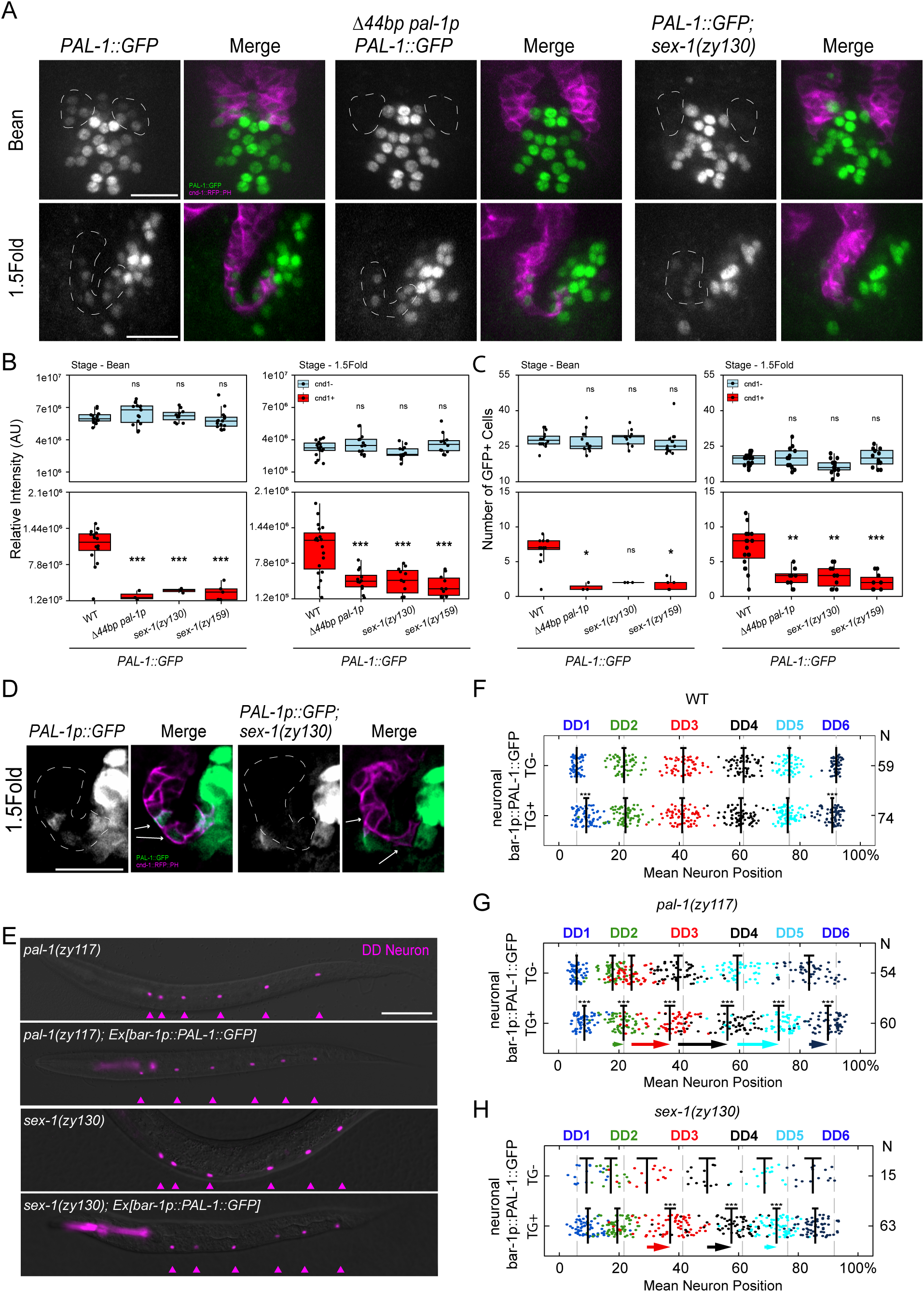
SEX-1 regulates PAL-1 expression. (A) Representative maximum projection images of *PAL-1::GFP* in WT, *pal-1(zy117) and sex-1* mutant embryos at bean and 1.5-fold stage. Hashed circles indicate regions with altered expression in mutant embryos. (B-C) Quantification of the (B) mean intensity and (C) number of nuclei expressing *PAL-1::GFP* in WT, *pal-1(zy117)* and *sex-1* mutant embryos. Nuclei were grouped based on their co-expression with *cnd-1^+^* cells, labelling the VNC. Means and 95% confidence intervals are indicated for each population with individual measurements shown. At the bean stage, N = 16, 15, 11 and 15, for WT, *pal-1(zy117), sex-1(zy130)* and *sex-1(zy159*), respectively. At the 1.5-fold stage, N = 19, 13, 13 and 12, for WT, *pal-1(zy117), sex-1(zy130)* and *sex-1(zy159*), respectively. (D) Representative images showing *pal-1p::GFP* transcriptional reporter in WT or *sex-1(zy130)* background. Number of embryos examined: N = 9 for WT and N = 1 for *sex-1(zy130)*. Arrows indicate regions of interest. (E) Representative images of DD neuron positions in *pal-1(zy117)* and *sex-1(zy130)* mutants with or without a *bar-1p::PAL-1::GFP* transgene. Arrowheads mark the position of DD neurons. Scale bar = 20μm. (F and G) Quantification of DD neuron position relative to the SABVL neuron and the rectum in L1 stage (F) WT, (G) *pal-1(zy117)*, and (H) *sex-1(zy130)* with or without the extrachromosomal transgene. Means and 95% confidence intervals are indicated for each neuron and animals scored (N) indicated on the right. Hashed lines indicate the position of WT DD neurons. Arrows indicate the size and direction of the mean neuron position shift in each transgene expressing neuron from the mean position of their non-transgene equivalent. Statistics: (B) one-way ANOVA with Dunnet’s Post-hoc Comparison, (C) Kruskal-Wallis Test with Dunn’s Post-hoc Comparisons, (F-G) Weltch’s T-Test. *p<0.05, **p<0.01, ***p<0.001.

If PAL-1 functions downstream of SEX-1, then expressing *pal-1* from a heterologous promoter active in VNC progenitors should bypass the requirement for *sex-1* and rescue the defects observed in *sex-1* mutants. To test this, we constructed a transgene in which the *pal-1* coding sequence is driven by the *bar-1* promoter, which is also active in DD and DA progenitors (Chan et al., 2025). This construct successfully rescued *zy117* defects and caused only minimal position defects when overexpressed in WT animals (Fig. 5E, F and G). When introduced into a *sex-1(zy130)* mutant background, the transgene was able to partially rescue neuronal positioning defects (Fig. 5E, F and H). These results are consistent with *pal-1* acting downstream of *sex-1* and, in agreement with our double mutant analysis, suggest that it is likely one of multiple targets through which SEX-1 mediates its effects on neuronal positioning.

## Discussion

The integration of multiple signaling pathways is key to proper nervous system development. *Caudal* homeobox (HOX) transcription factors, including the CDX family in mammals and PAL-1 in *C. elegans,* act as key mediators of signaling pathways and are widely required for morphogenesis and cell-fate decisions during embryogenesis (van den Akker et al., 2002; Wilson et al., 2003; Palmer et al., 2021; Zhao et al., 2022). This includes early neural development, which depends on gene expression that is activated by retinoic acid, WNT, and HOX transcription factors (Nordström et al., 2006; Skromne et al., 2007; Young et al., 2009; Sturgeon et al., 2011; Sanchez-Ferras et al., 2016). In this study, we identify PAL-1/Caudal as a key regulator of VNC morphogenesis and show that its function relies in part on SEX-1/NHR-mediated regulation of *pal-1* expression, but also involves SEX-1–independent mechanisms.

### Disruption of a DD/DA progenitor-specific *pal-1* promoter element causes neuron position defects

By examining endogenous *pal-1::GFP* expression, we found that *pal-1* is expressed in a subset of DD, DA, and DB progenitors during the bean to 1.5-fold stages of VNC development. The *pal-1* mutants used in this study, particularly the 44 bp *zy117* deletion, disrupt a promoter element that specifically drives expression in DD and DA progenitors but does not affect *pal-1* expression in DB progenitors. Although defects were observed in all three motor neuron classes (DD, DA, and DB), loss of *pal-1* expression driven by this promoter element primarily resulted in anteriorly mispositioned DD cell bodies, without apparent effect on embryonic or larval viability. In contrast, complete loss of PAL-1 activity results in embryonic and larval lethality due to defects in cell patterning and cell movement during embryogenesis that lead to severe morphological abnormalities (Edgar et al., 2001). Normal expression of cell fate markers and normal numbers of DD, DA, and DB neurons indicated that these defects were not due to alterations in cell fate.

*Cdx* genes are expressed in a temporally and spatially dynamic manner during early stages of mouse development. Gene expression begins in the primitive streak and tail bud early after implantation, then expands to neuroectodermal and mesodermal tissues, and gradually declines by late gestation (Meyer and Gruss, 1993; Beck et al., 1995; Chawengsaksophak et al., 1997; Benahmed et al., 2008). *pal-1* is similarly regulated in a complex temporal and spatial manner across multiple cell lineages (Hunter and Kenyon, 1996; Mootz et al., 2004; Edgar et al., 2001; Mainpal et al., 2011; Gilbert et al., 2020). This tightly controlled expression is mediated by multiple regulatory regions, including a fragment of exon 5 that drives *pal-1* transcription specifically in ventral hypodermal cells (Gilbert et al., 2020). Our work adds to this regulatory complexity by identifying a *pal-1* promoter element specific to DD/DA progenitors, suggesting that precise spatial control of *pal-1* expression in the developing VNC is critical for its role in neuronal cell body positioning.

### *pal-1/Cdx*, *vang-1/Vangl*, and *sax-3/Robo* act independently to regulate neuron position

We previously proposed that similarities in the cellular and molecular mechanisms underlying VNC morphogenesis and vertebrate neural tube formation might reflect an evolutionarily conserved process (Shah et al., 2017). Both involve convergent extension movements that narrow tissue along one axis while elongating it along another. They also share the formation and resolution of multicellular rosettes, the polarized distribution of PCP components such as VANG at cell membranes, and neighbour exchanges driven by actomyosin-mediated junctional contractions (Doudney et al., 2005; Ciruna et al., 2006; Ybot-Gonzalez et al., 2007; Williams et al., 2014). The involvement of the Caudal/CDX family protein PAL-1 in VNC assembly provides another point of similarity, as vertebrate CDX family members have also been implicated in PCP signaling during neurulation (Savory et al., 2011; Zhao et al., 2014). Consistent with this, loss of both *Cdx1* and *Cdx2* in mice results in the severe neural tube defect craniorachischisis (Savory et al., 2011; Zhao et al., 2014).

The neuronal cell body position defects in *pal-1(zy117)* are similar to those observed in *vang-1/Vangl* and *prkl-1/Prickle* mutants, which disrupt components of a PCP-like pathway, or in *sax-3/Robo* pathway mutants (Shah et al., 2017). These defects result from disrupted collective cell movements and failed intercalation of progenitor neurons at the midline during convergent extension, a process that brings progenitors to the midline and contributes to their single-file alignment and anterior-posterior distribution within the VNC. The *vang-1*/PCP-like and *sax-3/Robo* pathways function in independent, partially redundant pathways to regulate this process. Simultaneous disruption of both pathways severely impairs convergent extension, causing DD, DA, and DB progenitors to fail to move posteriorly, resulting in a clustering of neurons in the anterior VNC at hatching (Shah et al., 2017). A role for PAL-1 in these morphogenetic events would be consistent with its previously characterized functions. In particular, tissue-specific loss of *pal-1* in a subset of ventral hypodermal cells disrupts cell migration and intercalation during late embryogenesis (Gilbert et al., 2020), and *pal-1* is also required for the migration of several dorsal hypodermal cell populations (Edgar et al., 2001).

Both *pal-1(zy117)* and the null mutant *vang-1(tm1422)* exhibit anterior mispositioning of DD and DA cell bodies, although the displacement of DD neurons is significantly more severe in *zy117*. This occurs despite *zy117* not being a null allele, but rather disrupting *pal-1* expression in only a subset of DD and DA neurons. *vang-1; pal-1* double mutants exhibit a striking synergistic enhancement of neuronal position defects compared to either single mutant. This suggests that *pal-1* and *vang-1* act in parallel or independent pathways to regulate neuronal cell body position. This finding contrasts with the role of CDX proteins during neurulation, which involves PCP signaling through the protein tyrosine kinase PTK7 (Savory et al., 2011), a protein that lacks an identified homologue in *C. elegans*. However, since the developmental processes underlying VNC formation are not fully understood, and likely involve the interaction of several mechanisms, we cannot exclude the possibility that PAL-1 contributes to PCP regulation in some context, in addition to acting through independent pathways. A similar conclusion can be drawn from the observation that most DD and DA cell bodies are more severely displaced toward the anterior in *sax-3; pal-1* double mutants compared to either single mutant.

### *pal-1/Cdx* expression is regulated by SEX-1/NHR in DD and DA progenitors

The nuclear hormone receptor (NHR) SEX-1, which functions as a transcription factor, is primarily known for its role in sex determination and dosage compensation, acting in part through inhibition of the male-inducing transcription factor *xol-1* (Skipper et al., 1999; Gladden and Meyer, 2007; Meyer et al., 2010; Farboud et al., 2013). We found that SEX-1 is required for *pal-1* expression in DD and DA progenitors, representing a novel function for SEX-1 unrelated to sex determination. *sex-1* and *pal-1* mutants both exhibit anteriorly mispositioned DD and DA neuron cell bodies, although *pal-1* primarily affects DD neurons, whereas *sex-1* disrupts the positioning of both DD and DA neurons. The shared DD defects suggest that *sex-1* is important for DD neuron positioning, at least in part through its regulation of *pal-1* expression. However, it remains unclear whether this regulation involves direct binding of SEX-1 to the DD/DA promoter element in *pal-1*. The severe enhancement of DD and DA position defects in *sex-1(zy130); pal-1(zy117)* double mutants suggest that *sex-1* and *pal-1* also act in parallel or independent pathways to regulate neuron cell body positioning in the VNC. Since these defects resemble those caused by a strong disruption of convergent extension during early VNC formation, this suggests that both *sex-1* and *pal-1* contribute to convergent extension.

The relationship between SEX-1/NHR and PAL-1/Cdx in VNC formation parallels the roles of their homologues in morphogenetic processes involving tissue shape changes or convergent extension. CDX family members are regulated by both autonomous and non-autonomous signals, include the canonical WNT signaling pathway and NHRs (Houle et al., 2003; Sanchez-Ferras et al., 2012; reviewed in Bodofsky et al., 2017). Prominent NHR families like the retinoic acid receptors (RARs), retinoid X receptors (RXR) and the orphan NHR *Xenopus* germ cell nuclear factor (xGCNF) play key roles in the regulation of neurulation or cellular behaviors (Wendling et al., 1999; Mic et al., 2002; Barreto et al., 2003; Kam et al., 2013; Gur et al., 2022; Koch et al., 2025). This process may be partly achieved through the regulation of convergent extension, which is disrupted by genetic manipulation of retinoic acid signaling or manipulation of xGCNF (Barreto et al., 2003; Gur et al., 2022). Of these families, retinoic acid receptors, and their endogenous retinoic acid ligands, as well as thyroid receptors have also been shown to regulate *cdx* gene expression in several different contexts (Plateroti et al., 2001; Houle et al., 2003). For example, retinoic acid promotes and maintains *cdx* expression during the specification of multiple mesodermal cell populations, including in early cardiogenesis, zebrafish pronephron development, and axial extension during late gastrulation in mice (Houle et al., 2003; Wingert et al., 2007; Lengerke et al., 2012). Interestingly, SEX-1 shares some sequence similarity with the NHR1 superfamily, which includes retinoic acid receptors (RARs) (Carmi et al., 1998). Overall, the involvement of NHR and Cdx family members in VNC formation and neurulation points to a conserved set of cellular and molecular mechanisms underlying central nerve cord development.

## Materials and Methods

**Table 1:** Key Resources.

| Reagent or resource | Source | Identifier |
| --- | --- | --- |
| Chemicals, Peptides, and Recombinant Proteins |  |  |
| Levamisol hydrochloride | Millipore Sigma | 31742;<br>CAS:16595-80-5 |
| Bacterial and Virus Strains |  |  |
| *E. coli*: OP50 | Caenorhabditis Genetics Center (CGC) | RRID: WB-STRAIN:WBStrain00041969 |
| Experimental Models: Organisms/Strains |  |  |
| *C. elegans*: N2 (wild isolate) | CGC |  |
| *C. elegans*: OU792 (*unc-4*(*zy123*[*unc-4*::*mNG*])*zySi2*[*unc-30p*::*mcherry*::*H2B*::*unc-30*]; *vab-7*(*zy142*[*vab7*::*mNG*::*T2A*::*mScarlet-I*::*H2B*]) | Saharkhiz et al., 2024 | N/A |
| *C. elegans*: OU734 (*unc-4*(*zy123*[*unc-4*::*mNG*])*zySi2*[*unc-30p*::*mcherry*::*H2B*::*unc-30*]) | Saharkhiz et al., 2024 | N/A |
| *C. elegans*: *pal-1*(*zy43*) III | This study | N/A |
| *C. elegans*: *pal-1*(*zy44*) III | This study | N/A |
| *C. elegans*: *pal-1*(*zy117*) III | This study | N/A |
| *C. elegans*: *sex-1*(*zy130*) X | This study | N/A |
| *C. elegans*: *sex-1*(*zy159*) X | This study | N/A |
| *C. elegans*: *pal-1*(*zy112*[*pal-1*::*GFP*]) III | This study | N/A |
| *C. elegans*: *pal-1*(*zy117*) *pal-1*(*zy122*[*pal-1*::*GFP*]) III | This study | N/A |
| *C. elegans*: *sex-1*(*zy133*[*zy133*[*sex-1*::*mNG*::*AID*]) X | This study | N/A |
| *C. elegans*: *ynIs37[flp-13p::gfp]* III | CGC | WB Strain:<br>NY2037;<br>WormBase:<br>WBStrain00029158 |
| *C. elegans*: *sax-3(zy5)* X | Shah et al., 2017 | N/A |
| *C. elegans*: *vang-1(tm1422)* X | Mitani, 2017 | WB Strain:<br>RB1125;<br>WormBase:<br>WBStrain00031829 |
| *C. elegans*: *zyIs36(cnd-1::PH::mCherry; myo-2p::mCherry)* X | Shah et al., 2017 | N/A |
| Oligonucleotides |  |  |
| DNA Oligonucleotides | Eurofins | See Table S1 for sequences and use. |
| Software and Algorithms |  |  |
| Zeiss Zen 3.2 | Zeiss | https://www.zeiss.com/microscopy/en/products/software/zeiss-zen.html |
| R 4.4.2 | R Core Team, 2019 | N/A |
| ggplot2 | Wilkinson, 2011 | https://github.com/tidyverse/ggplot2 |
| Imaris 10 | Oxford Instruments | https://imaris.oxinst.com/ |

### Strains and Maintenance

All strains were maintained and examined at 20 C. The Bristol N2 strain was used as wild type (WT), along with the following alleles and transgenes: LGII: *unc-4(syb1658[unc-4::GFP]).* LGIII: *pal-1(zy43), pal-1(zy44), pal-1(zy117)*. LGX: *sex-1(zy45), sex-1(zy130), sex-1(zy159), sax-3(zy5), vang-1(tm1422), zyIs36[cnd-1p::PH::mCherry myo-2p::mCherry]*.

### pal-1 and sex-1 alleles

*pal-1(zy43), pal-1(zy44)* and *sex-1(zy45)* were identified in a genetic screen for DD neuron position defects (A. Colavita, unpublished). Whole-genome sequencing, as described in Noblett et al. (2019), was used to identify the genes and their associated DNA lesions. Mutants were subsequently outcrossed at least three times and reverified by Sanger sequencing (performed at OHRI StemCore) using primers N35-N37 for *pal-1* (all primers listed in Supplemental Table 1).

The *pal-1(zy117)* deletion was generated via DNA repair of CRISPR-Cas9-induced double strand breaks (Waaijers et al., 2013). Briefly, the target site (GAATGTCATAATGACCTTTT) was identified using the IDT guide RNA design tool at idtdna.com. The sequence was inserted into the Cas9-sgRNA plasmid pDD162 (Addgene #47549) using the NEB Q5 Site-Directed Mutagenesis Kit to generate plasmid pAC564. Young adult worms containing the DD reporter transgene *ynIs37* were injected with a mix of 50 ng/µl of pAC564, 2.5 ng/µl of myo-2p::mCherry (pCFJ90, Addgene #19327) and pBluescript plasmid (Stratagene) to a concentration of 100 ng/µl. Progeny were screened for DD position defects that resembled those in *zy43* and *zy44*. Sequencing (OHRI StemCore) with primers N35 and N36 was used to identify the deletion. Subsequent genotyping was performed using primers N35-N37.

### CRISPR-Cas9-mediated knock-ins

A GFP::3xFLAG and an mNG::AID::3xFLAG were inserted at the C-terminus of the endogenous *pal-1* and *sex-1* loci, respectively, using the CRISPR-Cas9 self-excising drug selection cassette approach described in Dickinson et al. (2015). Target sequences for *pal-1* (GTCGTGACAAACAGAAGATT) and *sex-1* (CAAAGGTGGCTAGCAGTGAA) insertions were cloned into the Cas9–sgRNA plasmid pDD162 to generate plasmids pAC487 and pAC703, respectively. The homology directed repair plasmids for *pal-1* (pAC486) and *sex-1* (pAC722) were constructed using the GFP (pDD282, Addgene #66823) and mNG::AID (pJW1582, Addgene #154312) selection cassette plasmids, respectively. The resulting knock-in alleles are *pal-1(zy112[pal-1::GFP])* and *sex-1(zy133[sex-1::mNG::AID])*.

### Reporter and rescue constructs

A *pal-1p::mNG* transcriptional reporter (pAC499) was constructed using Gibson Assembly (New England Biolabs). This reporter combines approximately 1.8 kb of distal and 2.4 kb of proximal promoter elements upstream of an mNG cassette, with the pPD95.77 vector (a gift from Andrew Fire, Stanford) serving as the backbone. Primers N48-N53 were used for the assembly (Table S1).

A promoterless *pal-1::GFP* minigene plasmid (pAC936) was constructed using Gibson Assembly. This construct contains the *pal-1* genomic coding region, with introns 3 and 5 removed, fused to the *GFP::3X Flag* cassette from the endogenous knock-in allele *pal-1*(zy112). Primers N54-N61 were used for the assembly (Table S1). The *bar-1p::pal-1::GFP* rescue construct (pAC937) was made by inserting a 4.9 kb *bar-1* promoter into pAC936 using Gibson Assembly with primers N62-N65 (Table S1).

pAC499 and pAC936 were each injected at a concentration of 5 ng/µl, along with 2.5 ng/µl of *myo-2p::mCherry* (pCFJ90) and 92.5 ng/µl of pBluescript plasmid.

### Embryonic Imaging

Widefield imaging was used to measure VNC rosette lifetimes. For widefield imaging, embryos were prepared as in Bao and Murray (2011). Briefly, several embryos were transferred from plates using a capillary attached to a glass pipette and mounted in a 3 µL drop of M9 containing approximately fifty 25 µm diameter polystyrene beads (Polyscience Inc.). The preparation was then sealed under a No. 1.5 coverslip using Vaseline. All widefield imaging was conducted on a Zeiss Imager M2, equipped with a Zeiss SVB-1 microscope signal distribution box/*TTL* trigger cable-controlled camera shutter to reduce phototoxicity. Single time-point images were taken across 15-30 0.5 µm slices at 63x (1.4NA). Unless otherwise stated, all images were deconvolved prior to analysis using constrained iterative deconvolution on Zen 3.2 (Zeiss) with a Gaussian algorithm, corrections for background and saturated pixels.

### Quantification of motor neuron position defects in newly hatched larvae

Quantification of neuron position defects in L1 larvae was performed as follows. First stage larvae (L1) were mounted on 2% agarose pads using 200 µM levamisole (#31742, Sigma). Images were acquired using a Zeiss Imager M2, Colibri7 LED source, AxiocamHR camera and taken at 40x (1.4NA) over 5-7 0.5um slices. Brightfield, RFP and GFP channels were acquired. Images were measured on FIJI/Image J software, version 2.14. Using the segmented line tool, the position of each DD, DA and DB neuron (or only DD and DA neurons in OU734) was measured along the worm. These positions were then converted into percentage locations using SABVR = 0% and the anus = 100%. Each percentage location was plotted using base R, version 4.4.2 (R Core Team, 2019) and geom_jitter from the ggplot2 suite (Wilkinson, 2011).

### Quantification of PAL-1 expression

Embryonic imaging of *pal-1(zy112[pal-1::GFP])* and *sex-1(zy133[sex-1::mNG::AID])* was conducted on a DM16000B Quorum Spinning Disk (Leica) with a EMCCD (Hamamatsu) camera. Single time-point images were taken at 63x (1.4NA) with a pinhole size of 50um. Images were acquired with a Brightfield exposure of 223ms, at FITC exposure of 500ms and a Cy3 exposure of 700ms, across 12 0.5um slices.

Intensity measurements of *pal-1(zy112[pal-1::GFP])* were conducted using Imaris 10 (Bitplane) after data was processed using Fiji/ImageJ. Briefly, Z-stacks were opened in their native format in Fiji/ImageJ. Both red and green channels were merged to form a composite color image, which was converted into RGB before exporting as a TIFF. Data was then imported into Imaris. Semi-automated spot detection was used to identify GFP nuclei with a XY diameter of at least 1.15 µm and a quality above 42. Spots were classified using the Imaris machine learning tool based upon the RFP channel and their proximity to the RFP labelled VNC cells. After spot creation was complete, the original GFP image was used to remove false positive spots which were outside the embryo boundary. Spots details including class and SUM Intensity were exported for statistical analysis and graphing.

### Statistics

Tests and the number of samples for each dataset are indicated in individual figure legends. Parametric data was analyzed using Weltch’s T-Test or an analysis of variance (ANOVA), as appropriate. Proportional data was compared using a Chi-squared test with Monte Carlo simulation for 10,000 replicates. When this returned a significant result, a pairwise analysis was conducted using Fisher’s Exact Test with Monte Carlo simulation for 10,000 replicates, adjusted with Holm corrections for multiple comparisons. Statistics were conducted using base R, version 4.4.2 (R Core Team, 2019), and the “multicomp” package for Dunnett’s Multiple Comparison Test (Bretz et al., 2011). Measurements related to embryonic phenotypes (expression pattern and order measurements) were scored blind to genotype using the Blind Analysis Tools plugin for Fiji/ImageJ (https://imagej.net/plugins/blind-analysis-tools).

## Acknowledgements

We thank Dr. Shohei Mitani for providing *vang-1(tm1422)* and Dr. Chris Li for providing *ynIs37*. Some strains were provided by the *Caenorhabditis* Genetics Center, which is funded National Institutes of Health Office of Research Infrastructure Programs (P40 OD010440). Data for *C. elegans* allele sequences was accessed through WormBase (http://www.wormbase.org, release WS294). The authors acknowledge the Cell Biology and Image Acquisition Core (RRID: SCR_021845) funded by the University of Ottawa, Natural Sciences and Engineering Research Council of Canada, and the Canada Foundation for Innovation. We would also like to acknowledge the assistance of StemCore Laboratories Genomics Core Facility (OHRI), RRID:SCR_012601. Studies presented in this report were supported by a Project Grant from the Canadian Institutes of Health Research (CIHR 123513 and 156160) to A. Colavita.

## Supplemental Material

**Supplementary Table 1. Oligonucleotides used in this study.**

**Supplementary Figure 1. Distribution of all DD, DA, and DB positions used to plot mean positions in Figure 2**. (A-C) Quantification of (A) DA, (B) DB and (C) DD neuron position relative to the SABVL neuron and rectum in L1 stage worms. Data shows individual samples for the data presented in Figure 2. Neurons are plotted along the AP axis, where SABVL and the rectum mark the 0% and 100% positions respectively. Black bars represent the means and 95% confidence intervals.

**Supplementary Figure 2. Distribution of all DD and DA positions used to plot mean positions in Figure 3 and 4**. (A-B) Quantification of (A) DA and (B) DD neuron position relative to the SABVL neuron and rectum in L1 stage worms. Each graph shows individual samples for the data presented in Figures 3 and 4. Neurons are plotted along the AP axis, where SABVL and the rectum mark the 0% and 100% positions respectively. Black bars represent the means and 95% confidence intervals.

**Supplementary Figure 3. Comparison of *pal-1(zy117)* double mutants with *vang-1(tm1422)* and *sax-3(zy5)*.** (A-B) Each graph shows the mean position of DD and DA neurons in larvae as presented in Figure 3. Neurons are plotted along the AP axis, where SABVL and rectum mark the 0% and 100% positions respectively. Means and 95% confidence intervals are indicated for each neuron and animals scored (N) indicated on the right. Hashed lines indicate the mean position of WT DD neurons. Arrows indicate the size and direction of the mean neuron position shift in the double mutant from the mean neuron position in (A) *pal-1(zy117)*, (B) *vang-1(tm1422)* or *sax-3(zy5)* single mutants. Statistics: one-way ANOVA with Tukey’s Post-hoc Comparisons. *p<0.05, **p<0.01, ***p<0.001.

**Supplementary Figure 4. Comparison of *sex-1(zy130); pal-1(zy117)* double mutants to single mutants.** (A-B) Each graph shows the mean position of DD and DA neurons in larvae as presented in Figure 4. Neurons are plotted along the AP axis, where SABVL and the rectum mark the 0% and 100% positions respectively. Means and 95% confidence intervals are indicated for each neuron and animals scored (N) indicated on the right. Hashed lines indicate the mean position of WT DD neurons. Arrows indicate the size and direction of the mean neuron position shift in the double mutant from the mean neuron position in (A) *pal-1(zy117)* or (B) *sex-1(zy130)* single mutants. Statistics: one-way ANOVA with Tukey’s Post-hoc Comparisons. *p<0.05, **p<0.01, ***p<0.001.

**Supplementary Figure 5. Loss of PAL-1 does not affect the expression of endogenous SEX-1::GFP.** Representative maximum projections of an endogenous *SEX-1::mNG* in WT (A) and *pal-1(zy117)* embryos (B). At the bean stage, N = 15 for both WT and *pal-1(zy117)*. At the 1.5-fold stage, N = 14 for WT and N = 15 for *pal-1(zy117)*. No visual changes are observed. Posterior *cnd-1p::RFP::PH* expression marks DD and DA neurons. The white hashed outline marks the embryo. Scale bar = 10μm.

