## Supplemental Table 1 for "Nuclear hormone receptor regulation of PAL-1/Caudal mediates ventral nerve cord assembly in *C. elegans*"

### *Supplemental Table 1. Oligonucleotide sequences used in this study.*

| Oligo # | DNA Oligos Sequence (5' – 3') | Source | Use |
|---|---|---|---|
| N34 | GAATGTCATAATGACCTTTTGTTTAAGAGCTATGCTGGAA | Eurofins | SGRNA for the deletion found in *pal-1(zy117)* |
| N35 | ACAACCTCAAGTACATACTGC | Eurofins | Sequence *pal-1(zy117)* |
| N36 | ACTACAACGATGTTCACATCC | Eurofins | Sequence *pal-1(zy117)* |
| N37 | CAATAATCTGACAGATGGTTGG | Eurofins | Sequence *pal-1(zy117)* |
| N38 | GTCGTGACAAACAGAAGATTGTTTAAGAGCTATGCTGGAA | Eurofins | SGRNA for *PAL-1::GFP* |
| N39 | ACGTTGTAAAACGACGGCCAGTCGCCGGCACACTCGCATGTTAAAGGGAG | Eurofins | Repair template for *PAL-1::GFP* |
| N40 | CATCGATGCTCCTGAGGCTCCCGATGCTCCTAGACGAATCTTCTGTTTGTCACGA | Eurofins | Repair template for *PAL-1::GFP* |
| N41 | CGTGATTACAAGGATGACGATGACAAGAGATAAGTACTCATCTACTTACAAAGAAAATGTAC | Eurofins | Repair template for *PAL-1::GFP* |
| N42 | GGAAACAGCTATGACCATGTTATCGATTTCGCAACTACCAGCGACTTAC | Eurofins | Repair template for *PAL-1::GFP* |
| N43 | CAAAGGTGGCTAGCAGTGAAGTTTAAGAGCTATGCTGGAA | Eurofins | SGRNA for the *SEX-1::mNG::AID* |
| N44 | ACGTTGTAAAACGACGGCCAGTCGCCGGCATATGGATCATTGGCGGACGA | Eurofins | Repair template for *SEX-1::mNG::AID* |
| N45 | CATCGATGCTCCTGAGGCTCCCGATGCTCCGCAGTGAACGGGAGTCTGTT | Eurofins | Repair template for *SEX-1::mNG::AID* |
| N46 | CGTGATTACAAGGATGACGATGACAAGAGATAGCCACCTTTGTCAGATTCT | Eurofins | Repair template for *SEX-1::mNG::AID* |
| N47 | GGAAACAGCTATGACCATGTTATCGATTTCTTCTGATAGAGGTCGACGG | Eurofins | Repair template for *SEX-1::mNG::AID* |
| N48 | ACAACTTGGAAATGAAATACTGCTTGAGATGCAAAGCGA | Eurofins | *pal-1p::mNG* fragment 1 |
| N49 | CGATCAGAGCTATCGAGTAC | Eurofins | *pal-1p::mNG* fragment 1 |
| N50 | GTACTCGATAGCTCTGATCGGTCAAGTCTGCCGTTGTTC | Eurofins | *pal-1p::mNG* fragment 2 |
| N51 | TCTAGAGTCGACCTGCAGTGGTTGCAGCATCTTTTCTCTG | Eurofins | *pal-1p::mNG* fragment 2 |
| N52 | CTGCAGGTCGACTCTAGA | Eurofins | *pal-1p::mNG* fragment 3 |
| N53 | TATTTCATTTCCAAGTTGT | Eurofins | *pal-1p::mNG* fragment 3 |
| N54 | ACAACGATGGATACGCTAACATGTCGGTCGATGTCAAG | Eurofins | promoterless *pal-1::GFP* fragment 1 |
| N55 | ATCCCTGAAACTGTTGATAATCCATAAACGG | Eurofins | promoterless *pal-1::GFP* fragment 1 |
| N56 | TTATCAACAGTTTCAGGGATTCTCGTGG | Eurofins | promoterless *pal-1::GFP* fragment 2 |
| N57 | CTTTGCACGCCTATTTTGAAACCAAATCTTGATTTG | Eurofins | promoterless *pal-1::GFP* fragment 2 |
| N58 | TTCAAAATAGGCGTGCAAAGGATCGTCG | Eurofins | promoterless *pal-1::GFP* fragment 3 |
| N59 | CCGACTAGTGGGCAGATCTTACTGGATAGTTAATCTCATCAATTTCTGC | Eurofins | promoterless *pal-1::GFP* fragment 3 |
| N60 | AAGATCTGCCCACTAGTC | Eurofins | promoterless *pal-1::GFP* fragment 4 |
| N61 | GTTAGCGTATCCATCGTTG | Eurofins | promoterless *pal-1::GFP* fragment 4 |
| N62 | ATACGCTAACATCGAGTTGAATCCTCTGGC | Eurofins | *bar-1p::pal-1::GFP* fragment 1 |
| N63 | CATCGACCGAGATCCAGGCCATCCCAGTTTTC | Eurofins | *bar-1p::pal-1::GFP* fragment 1 |
| N64 | GGCCTGGATCTCGGTCGATGTCAAGTCG | Eurofins | *bar-1p::pal-1::GFP* fragment 2 |
| N65 | TCAACTCGATGTTAGCGTATCCATCGTTG | Eurofins | *bar-1p::pal-1::GFP* fragment 2 |
