## Supplementary figures and images for "Nuclear hormone receptor regulation of PAL-1/Caudal mediates ventral nerve cord assembly in *C. elegans*"

### Supplemental Figures S1-S5

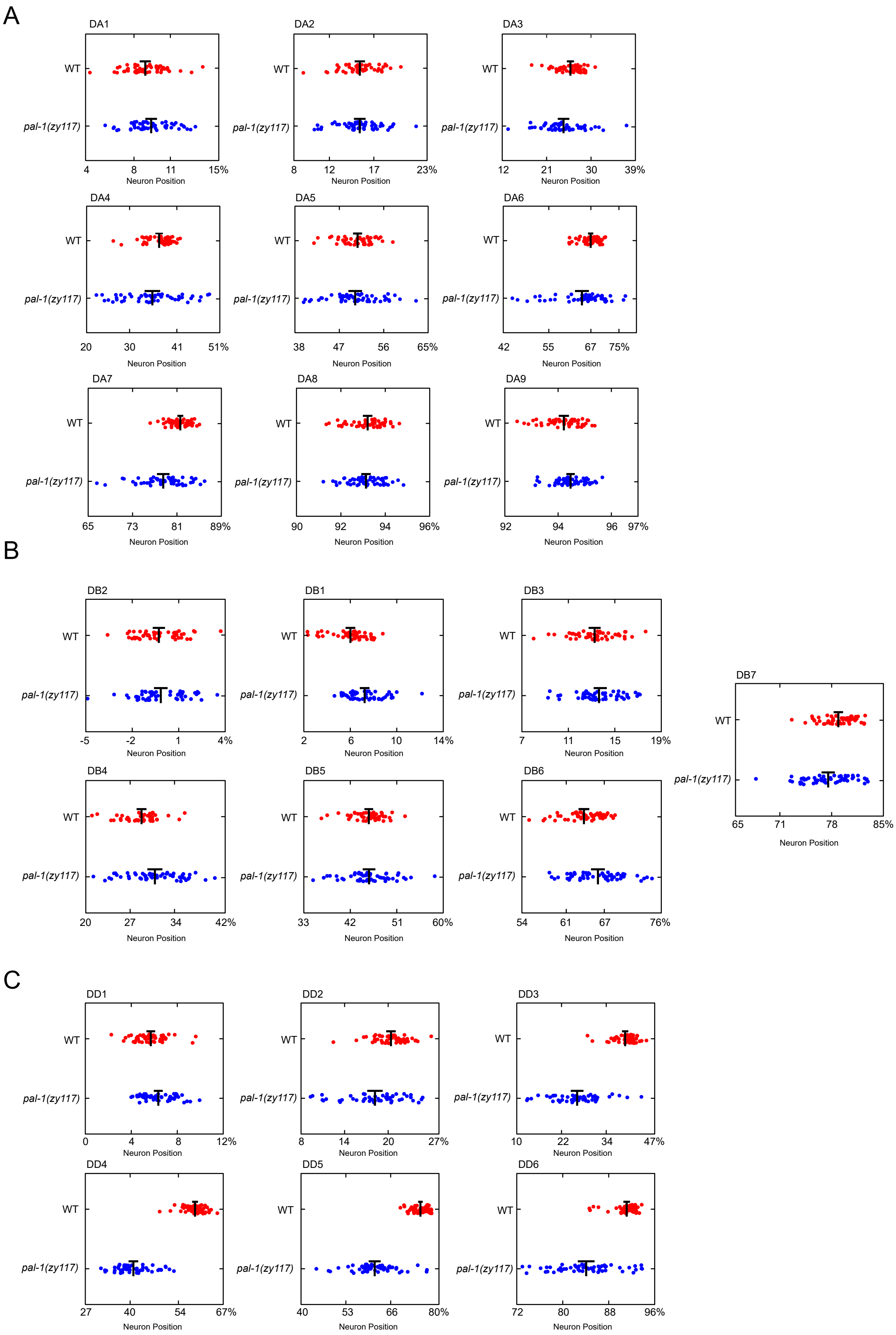

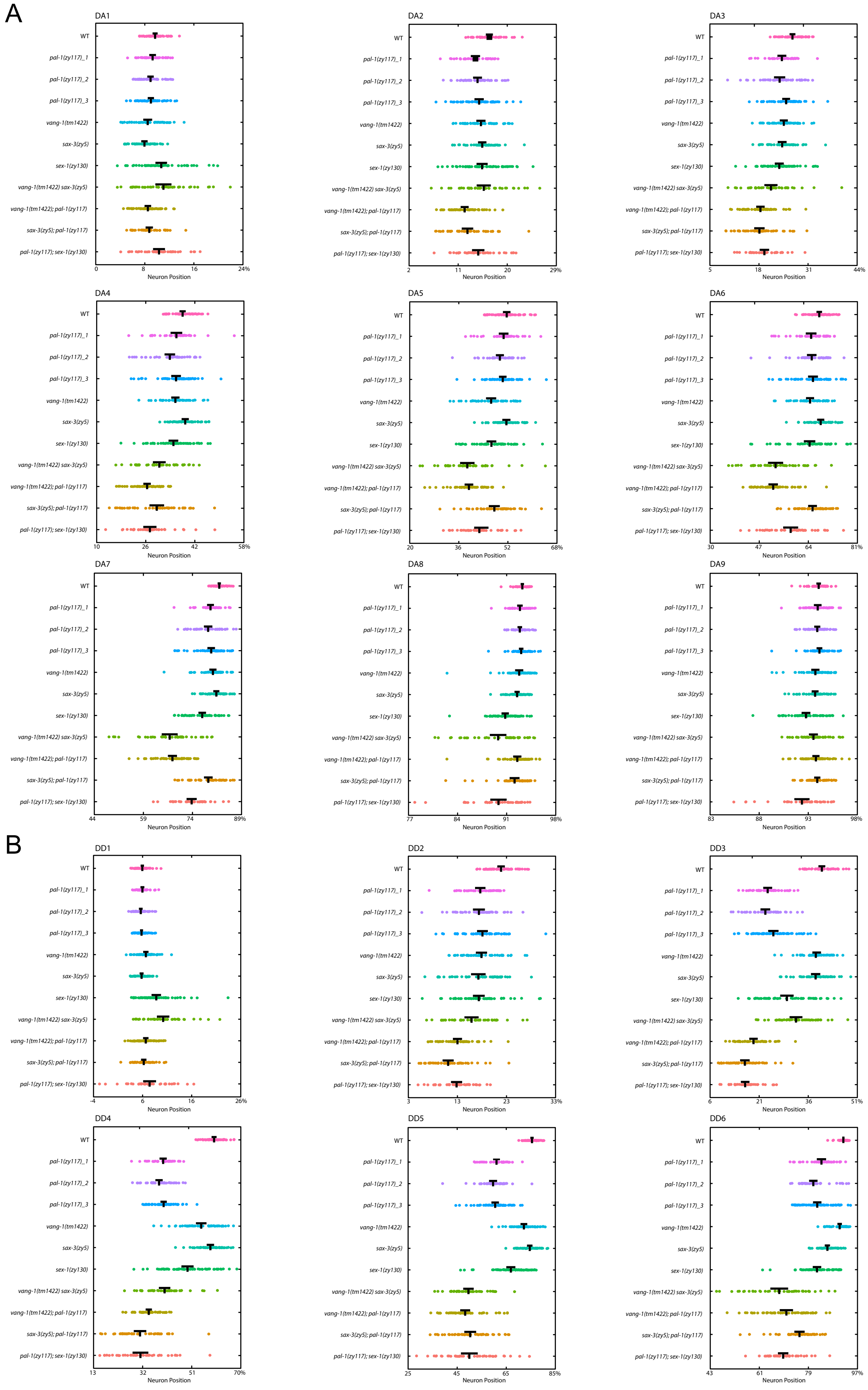

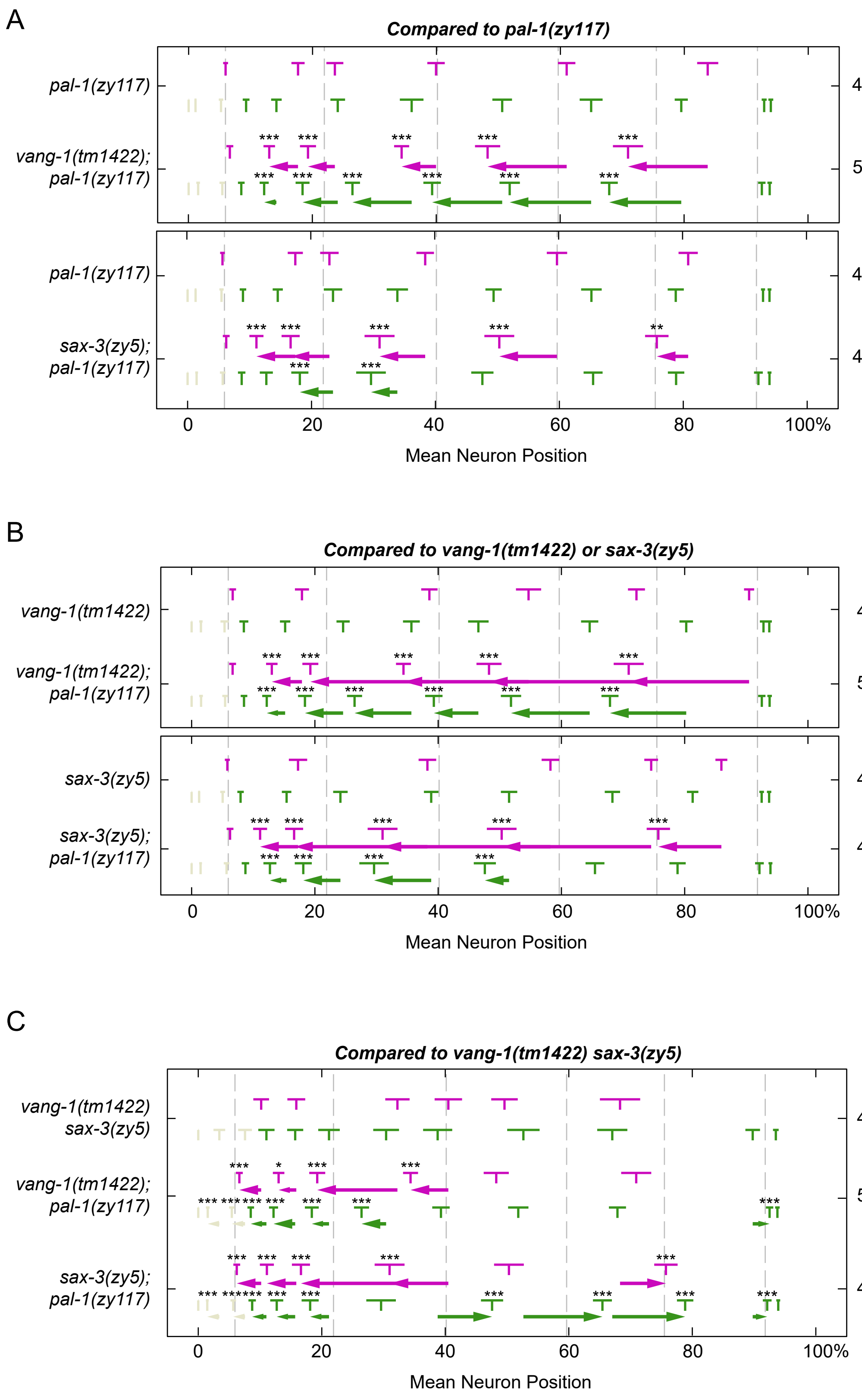

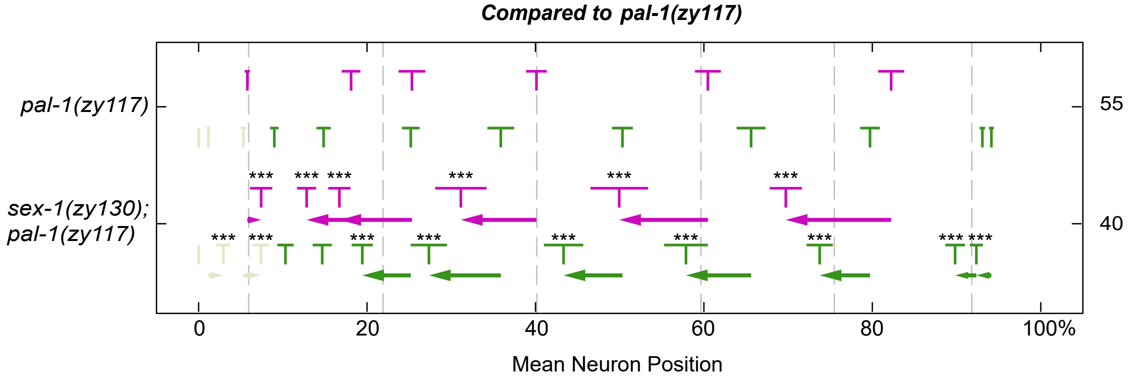

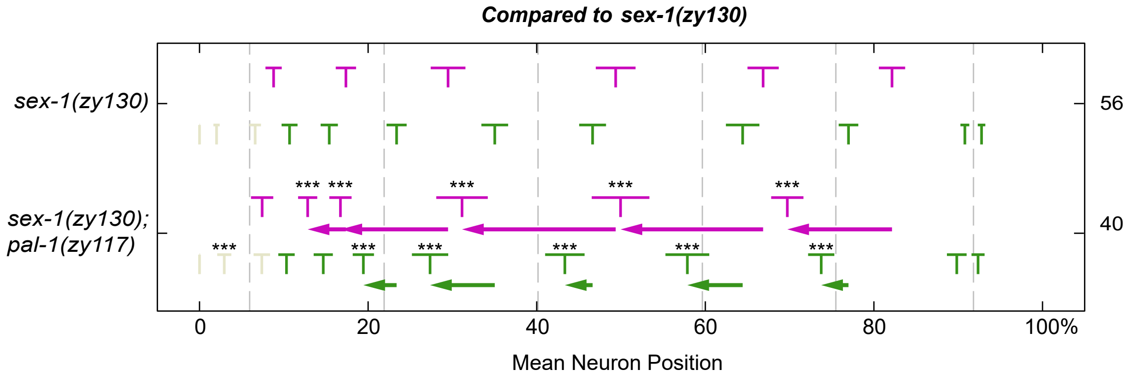

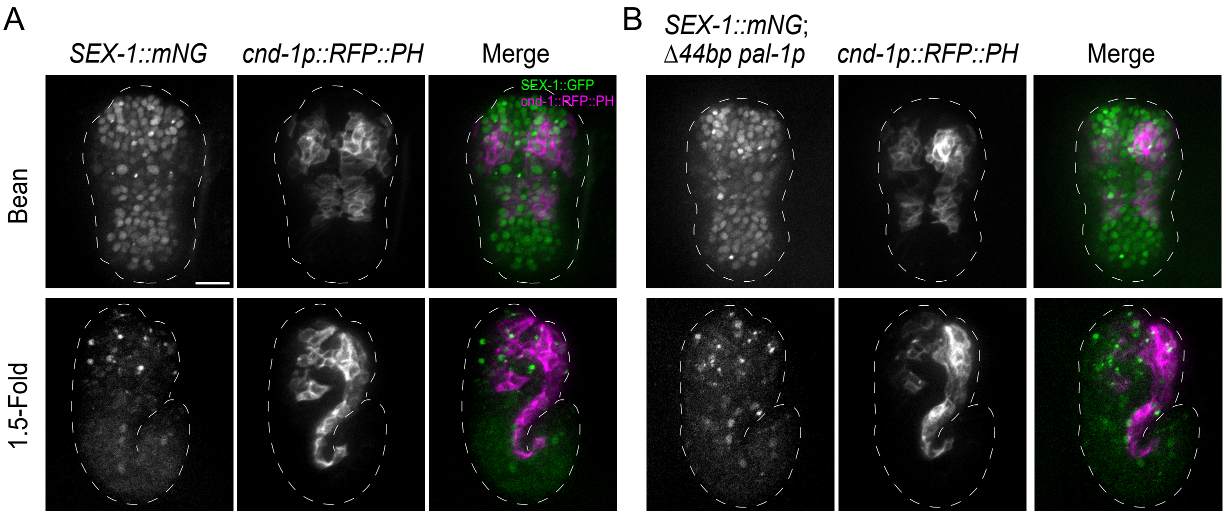
